# Filtering Methods to Improve the Accuracy of Indoor Positioning Data for Dairy Cows

**DOI:** 10.1101/186353

**Authors:** Matti Pastell, Lilli Frondelius, Mikko Järvinen, Juha Backman

## Abstract

Several indoor positioning systems for livestock buildings have been developed to be used as a tool in automated animal welfare monitoring. In many environments the measurements from positioning systems still contain unwanted noise and the quality of the measurement data can be enhanced using filters.

The aim of this study was to develop an efficient filter for positioning data measured from dairy cows with UWB-based indoor positioning system in a free stall barn. We developed and tested a heuristic jump filter combined with median filter and extended Kalman filter. The performance of the filters were compared against reference data collected from Insentec Roughage intake feeders and scan sampling of animal presence in a specific lying stall with over 1500 reference observations from both methods.

We were able to improve the quality of the positioning data significantly using filtering. The 9^th^ order median filter provided best estimates for cow position when the cows were not moving with median 100% of measurements located in correct stall and 84% in correct feeding trough when compared to the reference observations and measurements. The extended Kalman filter also improved the positioning accuracy significantly when compared to raw data and provides better of estimates of the trajectory of moving cows.

## 1. Introduction

Several indoor positioning systems for livestock buildings have been introduced to the market during past years. These systems are an interesting tool in automated animal welfare monitoring if sufficient positioning accuracy and sampling rate can be reached. First studies analyzing the data from positioning systems have shown that the data can be used to measure feeding time (Oberschätzl et al. 2015, Tullo et al. 2016), the effect of hoof lesions on walking distance (Frondelius et al. 2015), and the effect of lameness, estrus (Homer et al. 2013, Veissier et al. 2017) on cattle behaviour.

One of the first published systems in dairy cows was based on radar technology (Gygax et al. 2007). The system reached accuracy below 1m, but the battery life of the system in continuous operation was only 24 hours. Barn environment also has certain challenges for indoor positioning: usually there are metal structures and other obstacles that may cause reflections of signal and artefacts (Gygax et al. 2007). However, other experiments have shown that Ultra Wideband (**UWB**)-based systems can achieve positioning error below 1 m after cleaning of outliers (Porto et al. 2014) with a battery life from months to years depending on sampling rate and battery size.

In many environments the measurements from positioning systems still contain unwanted noise and the quality of the measurement data can be enhanced using filters. The Kalman filter is nowadays de facto filter to be used to estimate processes that can be modelled using mathematical model. The filter was originally proposed by Kalman (1960) and the first implementation was in space navigation in Apollo-program by Schmidt (Smith et.al 1962). The first implementation was also actually Extended Kalman Filter because linearization of the state transition model using Taylor series was used. Kalman filter and its extensions have shown to be suitable in wide range of applications. In agriculture, it has been used extensively for example in navigation applications (Oksanen et al. 2005, Lenain et al 2005, Backman et al 2010). The Kalman filter is ideal method to combine multiple measurement sources to get better state estimates, for example to reduce measurement noise, or to estimate states that cannot be directly measure, for example wheel slippage in navigation (Backman et al 2013). With indoor positioning data the Kalman filter has been previously used e.g. to estimate the position of cows based on Bluetooth radios (Tøgersen et al. 2010) and to track humans using sensor fusion of UWB positioning and inertial measurement unit (IMU) (Pulido Herrera 2009).

A median filter is a robust nonlinear filter that replaces the values in data with the moving median of the filtered point and neighboring points. It has been shown to improve the accuracy of human activity recognition based on UWB positioning data (Piltaver et al. 2015).

The aim of this study was to develop an efficient filter for positioning data measured from dairy cows with UWB-based indoor positioning system in a free stall barn. We introduce here a several options for improving the quality of the data and compare the results against reference datasets.

## 2. Materials and Methods

### 2.1 Measurements

#### Indoor positioning system

Ubisense real time location system (Ubisense, Germany) is based on Ultra Wide Band (UWB) radio signals. Animals are monitored by means of a collar-mounted tag (Ubisense Series 7000 Industrial tag) that transmits UWB pulses of extremely short duration, and remote sensors (Ubisense Series 7000 IP Sensors) which enable location to be mapped by using Time-Difference-of-Arrival (TDoA) and Angle-of-Arrival (AoA) techniques. The positioning system was set up to cover an area of 22 x 25 m with six sensors and calibrated for the area according to the system manual. We achieved static accuracy of ± 30 cm in the same dairy barn with no cows present (Frondelius et al. 2014).

The tags were attached to the collar of the cows, so that the tag is positioned on the top of the neck and the counter weight of the collar under the neck to keep collar still (Figure 1). Position of the collar and the tag was checked regularly and corrected if tag had moved more than 5 cm from the intended place.

**Figure 1.**
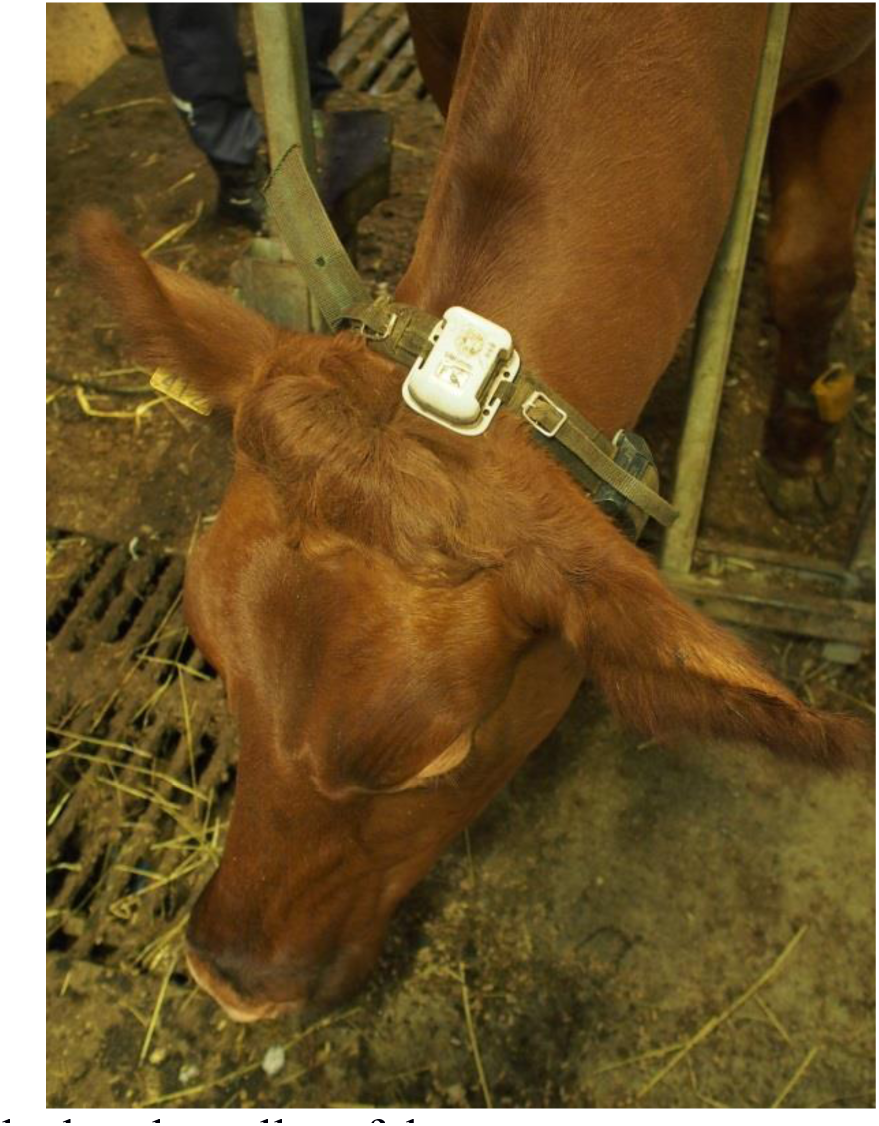
Ubisense tag attached to the collar of the cow.

#### Housing

Cows were loose-housed in a curtain-wall barn with rubber-matted stalls with steel stall separators and slatted passageways cleaned using manure robots in two sections of 24 cows. Both sections have their own concentrate feeder and 12 Insentec Roughage Intake Control system (RIC)-feeders with barrier structures preventing stealing behavior (Ruuska *et al.* 2014).

#### Reference data collection

We recorded the position data from 47 cows during 24 hours and collected simultaneously reference position data from Insentec RIC roughage intake measurement system. The experimental cows were 16 Nordic Red and 31 Holstein-Friesian lactating cows (21 primi-and 26 multiparous cows, 252 ± 211 (Mean±SD) days in milk, milk production 30.0 ± 6.3 kg/day (Mean±SD)). The RIC system is equipped with a RFID reader and photocell to identify when the cow is keeping the head in the trough. This resulted in total of 1668 recorded feeding bouts.

The cow position in a lying stall was recorded with direct observation scan sampling from 50 different cows from 47 of the 48 stalls in the barn during 4 days 6h/day resulting in total of 1510 observations.

The experimental cows were 17 Nordic Red and 33 Holstein-Friesian lactating cows (23 primi-and 27 multiparous cows, 241 ±182 (Mean±SD) days in milk, milk production 27.7±6.2 kg/day (Mean±SD)). During the observation the stall number where the animal was located and time were recorded. Only data from the cows that stayed still at least for 10 seconds during observation point were collected. The coordinates of the stalls and feeding troughs were recorded from the barn map calibrated to match the coordinates of the positioning system.

We compared the positioning accuracy and precision to the raw data and the after applying different filters. The results for raw and jump filtered data are presented without interpolation and median filters and extended Kalman filter (EKF) after interpolation.

### 2.2 Data analysis

Three different types of errors were identified from the measurement data and a strategy for correcting each was developed:

1. Large errors from reflection. These are characterized by large jumps of 2-15 m between samples and the tag returning to initial position after the jump. We developed a custom filter called jump filter for correcting these. The jump filter was always applied before other filters.
2. Normal errors. These are defined as random variation in position that can’t be clearly attributed to unrealistically large. These were filtered using either Extended Kalman Filter or Median filter after applying the jump filter and the performance of the methods was compared.
3. Missing data. The measurements from the positioning system in the used environment have quite a lot of missing data that needs to be interpolated before the data can be used to analyze animal behavior. We used interpolation to fill in missing samples after applying either the median filter or EKF.

#### Jump Filter

We developed a custom filter that we call the “jump filter” to correct large reflection errors from the data. The basic idea is to remove data points that clearly can’t be caused by cow movement.

We defined a criterion for omitting a data point p_t_ if the distance to previous p_t-1_ and next points p_t+1_ is more than 0.5m and p_t-1_ and p_t+1_ are close to each other:

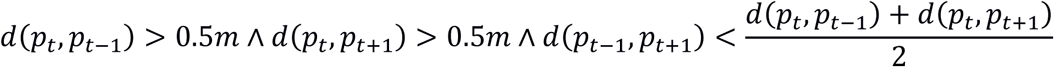

where the *d*(*p*_1_,*p*2) is the distance between two points. A schematic of the idea of removing errors is shown in Figure 2 and two examples of reflection errors before and after applying the filter are shown in Figure 3.

**Figure 2.**
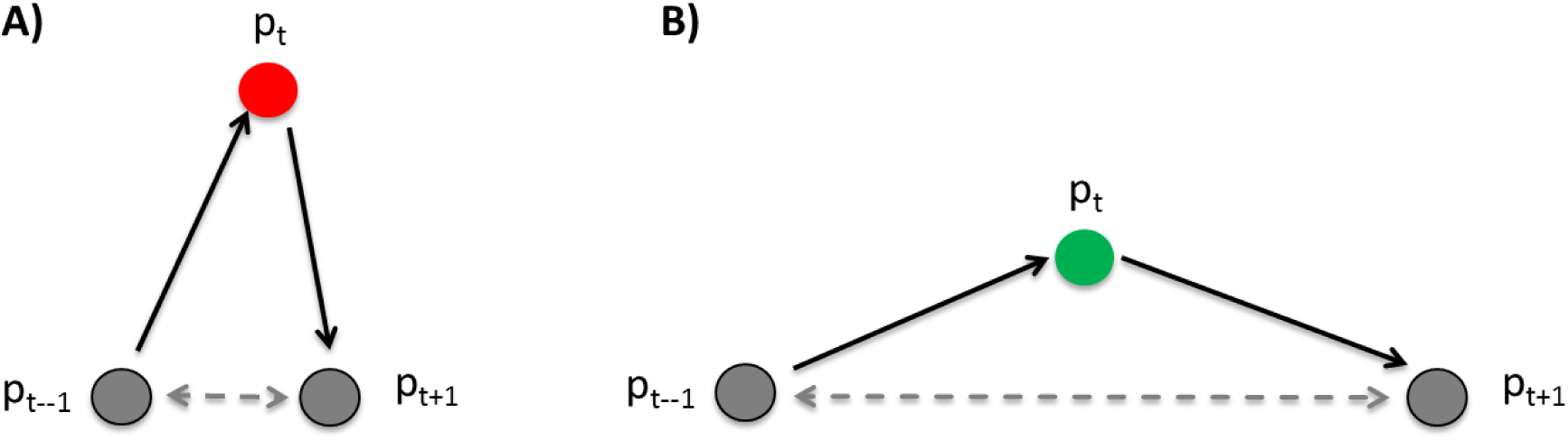
Example of the operation of the jump filter A) a sample makes a large jump and returns close to origin and is judged as erroneous B) a sample moves, but doesn’t return to origin and is judged as valid data point likely coming from a moving cow.

**Figure 3.**
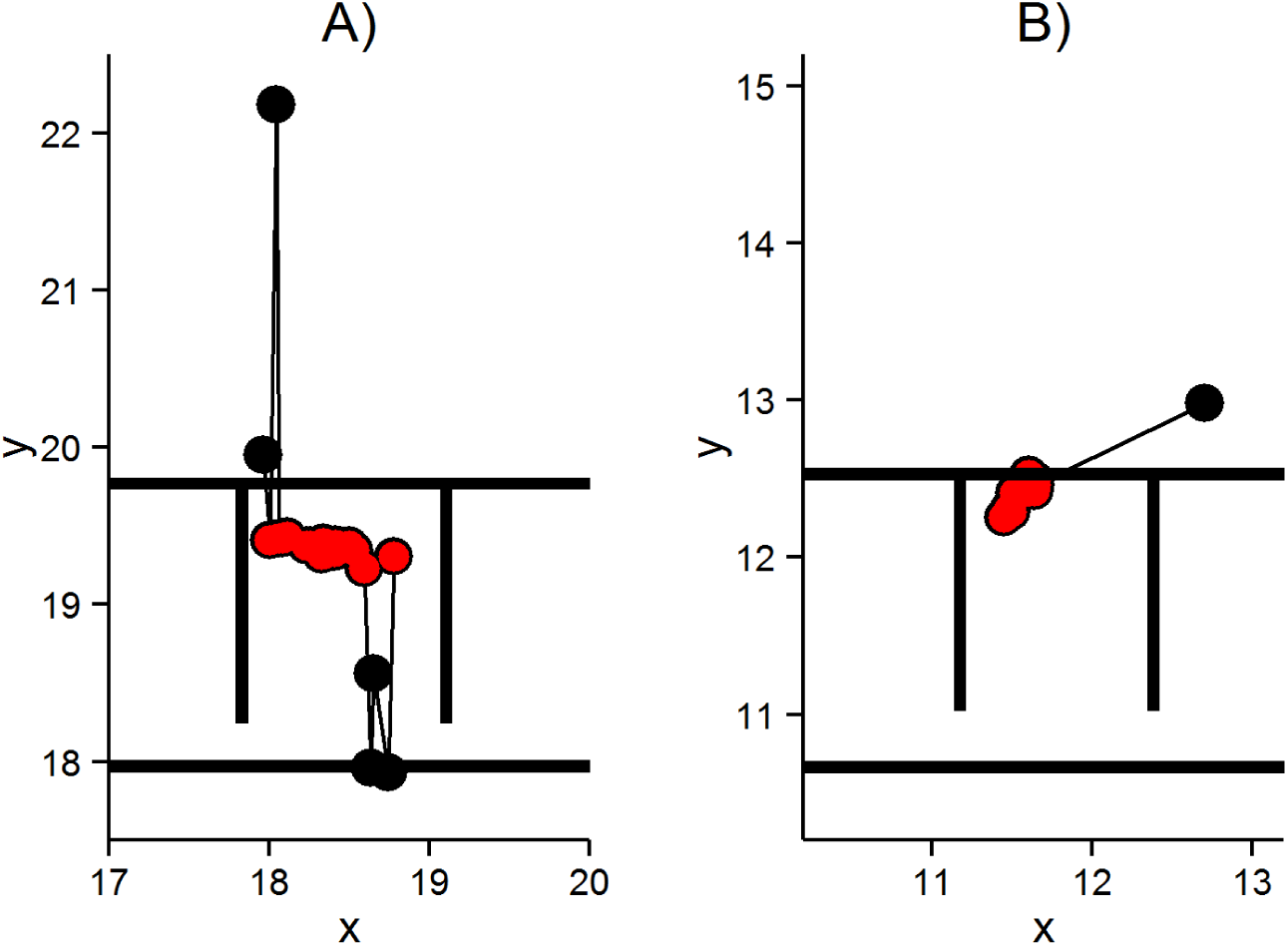
Two examples of jump filtered reflection errors when the cow is lying in stall. The black points represent raw data and red points data after applying the jump filter.

#### Extended Kalman Filter

We implemented EKF reduce random noise from the measurements. In this case the exact movement model of the cow is impossible to formulate and an approximation is used. The state of the cow is represented using the three dimensional position (x, y, z). In addition the movement is represented in the state vector using direction (θ) and speed (v) and corresponding time derivatives (ω and a):

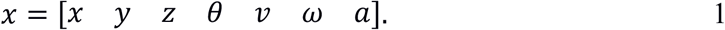

In the model, the position is modelled as a motion according to the speed and the direction of the movement. The direction and the speed are assumed to be random and modelled using modified Wiener process. The damping factor D_ω_ drives the rotational speed (ω) towards zero and the damping factors D_a_ and D_v_ drives the acceleration and speed towards zero respectively. The continuous time model is hence:

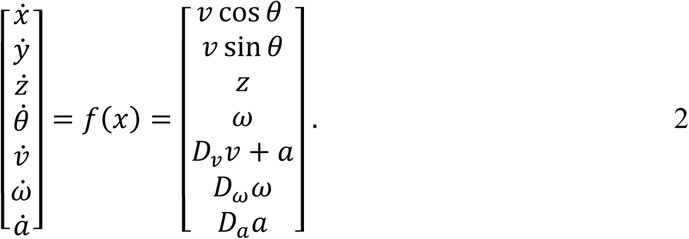

The damping factors (D_v_, D_a_ and D_w_) is calculated from the fall times (T_v_, T_a_ and T_w_) where the corresponding state has decreased to the one tenth of the original value:

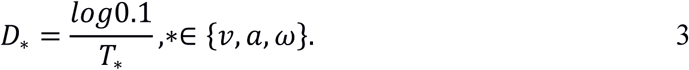

The cow position measurements are not synchronized and the time between consecutive measurements can vary between 0.8 seconds to several minutes. Therefore filtering is implemented using variable step length and hence the discretization time (*h*_*k*_) is not constant. For simplification we assumed that the speed (*v*_*k-1*_) and the direction (θ*k-1*) are constant, when position equations (*x*_*k*_ and *y*_*k*_) are discretized using Euler forward method. For the speed and direction equations (*θ*_*k*_, *v*_*k*_, ω_*k*_ and *a* _*k*_) and *a*_*k*_) exact solution is found. Discrete time model is hence:

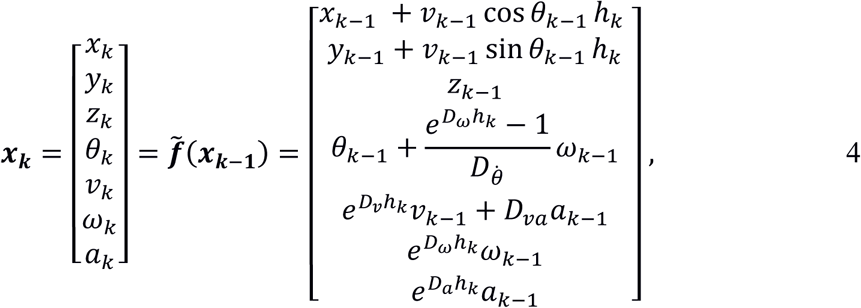

where

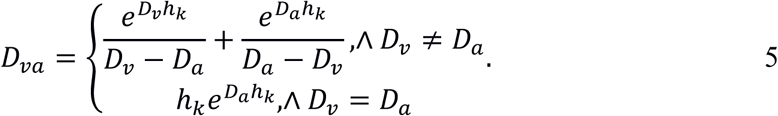

The Extended Kalman Filter also needs Jacobians of the discrete state transfer function (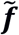) and the measurement function. The Jacobian of the state transfer function is:

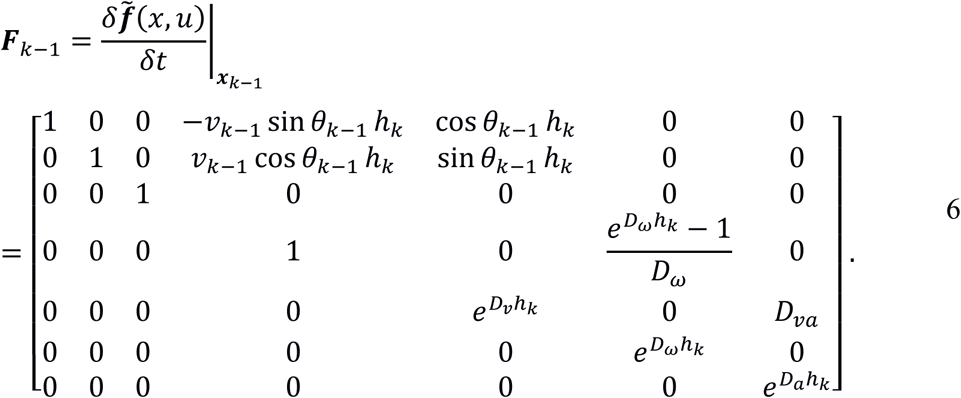

The measurement function is already linear because only the position is measured directly. Hence the corresponding Jacobian is:

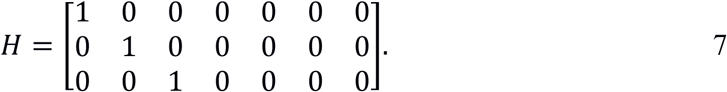

The resulting overall estimation model is:

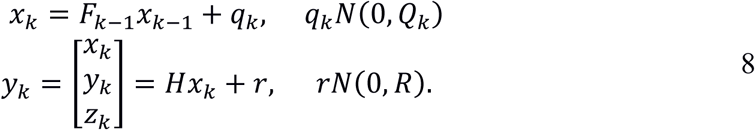

In the experiments, we manually tuned the parameters of the filter by trial and error method. The noise covariance matrices were:

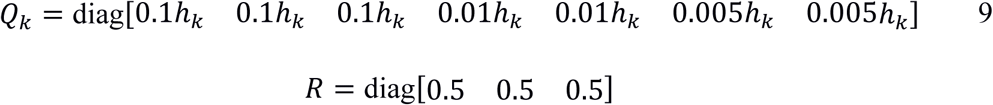

The falling times of the damping factors were:

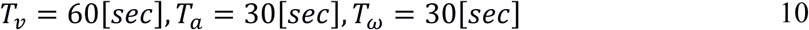

#### Median filter

We compared the EKF to median filters of different order. A median filter replaces the values in data with the moving median of the filtered point and neighboring points. The filter order ? specifies how many points are used to calculate the median. Higher filter order suppresses more noise, but underestimates the movement of the animal. The median filter is very robust against outliers and is well suited for finding the correct location when the animal is not moving.

#### Handling missing data

We chose to interpolate missing data using linear piecewise interpolation, because in most cases the tag reappeared close to the same location where the signal was lost. In addition the method is resistant against overshoot which caused problems with e.g. cubic interpolation even after applying the jump filter. We applied interpolation after EKF or median filtering.

#### Implementation

The data analysis was conducted using R 3.2.2 (R Core Team 2015) using RcppArmadillo (Eddelbuettel & Sanderson 2015) package and C++ for implementing the jump filter and extended Kalman filter. R function “runmed” using the Härdle & Steiger (1995) algorithm was used for the median filter. The ggplot2 (Wickham 2009) and beanplot (Kampstra 2008) packages were used for creating the plots.

## 3. Results

### 3.1 Missing data and jump filter

In average 58% of samples were missing during the trial. There was a large variation between number of received data points between cows, range 11% - 76% (Figure 4a). Most of the time missing data occurred in the stalls and near the edges of the positioning system coverage (Figure 5). Applying the jump filter further resulted in reduction (mean ± SD) of 9.8 ± 0.3% of the received samples. Milking times have been excluded from the analysis, because the milking parlour is located outside the sensor coverage.

**Figure 4.**
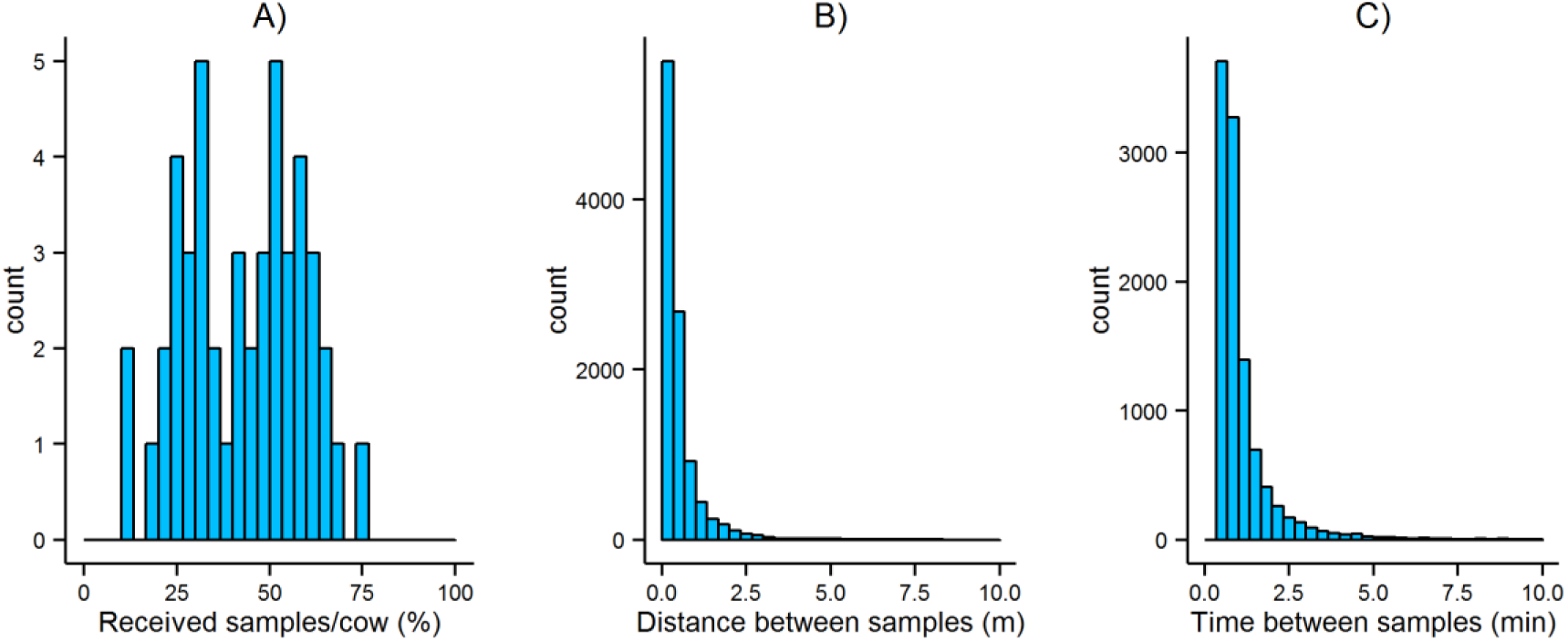
Analysis of missing data: A) Percentage of received samples/cow B) The distance the tags moved when the signal was lost during C) The distribution of the duration of signal loss. Both calculated from 24h sample from 47cows.

**Figure 5.**
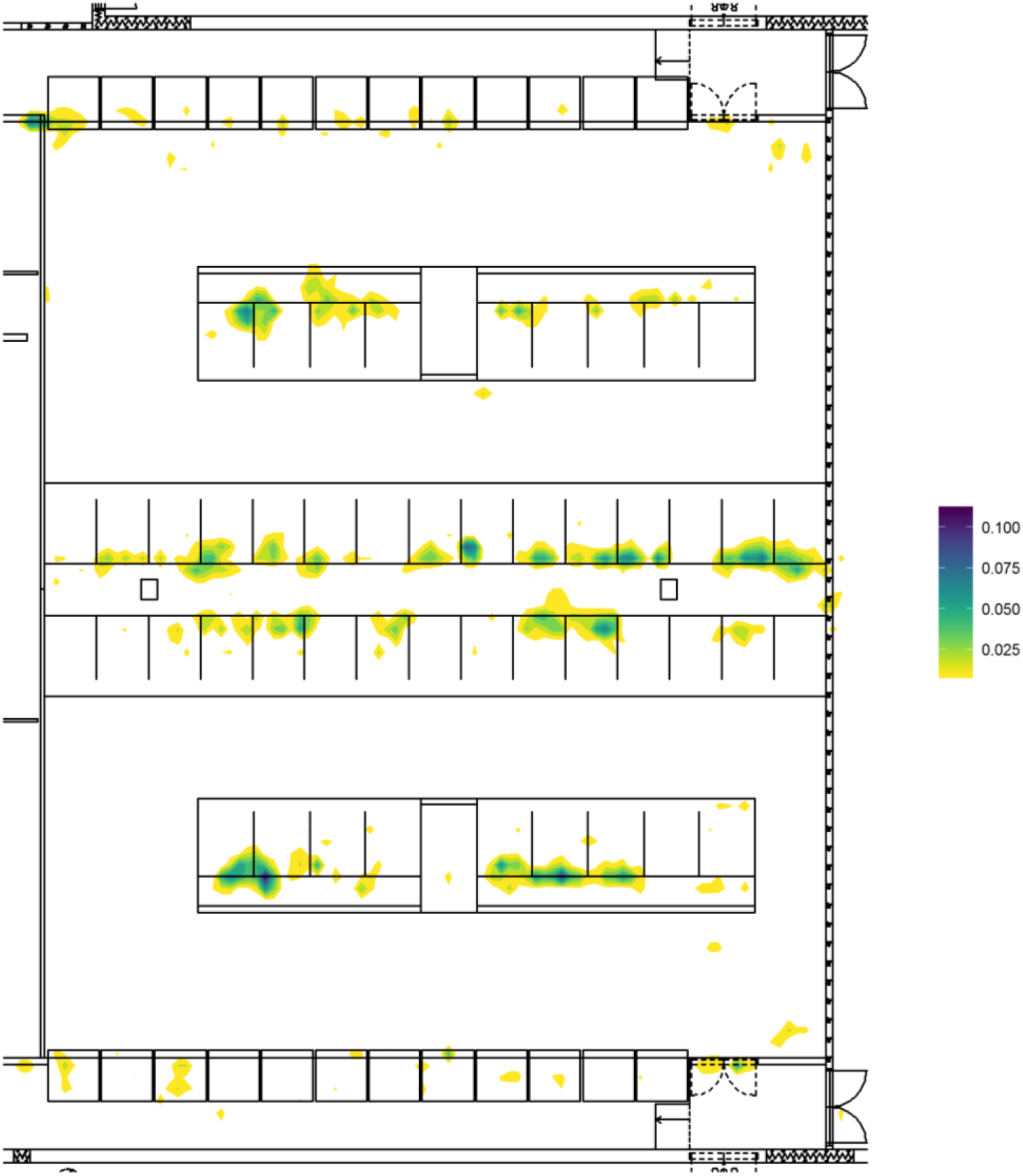
A density estimate of the areas were positioning signal was lost for more than 30s calculated from 24h sample from 47 cows.

The tags were lost for less than 1 minute in 66% of the cases and less than 2 minutes in 90% of the cases (Figure 4b) and the cows moved less than 1m in 88% of the cases during the time that we did not receive data from the tags (Figure 4c). This means in practice that the data can still be used to analyze the movement of the cows, because we receive most of the data when the cows are walking on the alleys.

### 3.2. The effect of filtering on positioning accuracy and precision

We calculated the position of the cows from the positioning data from a 30s bout and compared it to the manually scored stall location and we compared the positioning data during feed bouts to automatically recorded Insentec trough visit data. We calculated the proportion of the positioning measurements that located inside the correct stall and trough for each observation. The proportion of points located in the correct area as compared to reference data clearly increased with filtering (Figure 6) with best results achieved using 9pt median filter. For stalls the median proportion of points/observation hitting the correct stall increased from 86% to 100% and for feeding troughs from 53% to 84%. The median filter and EKF were applied to jump filtered data.

**Figure 6.**
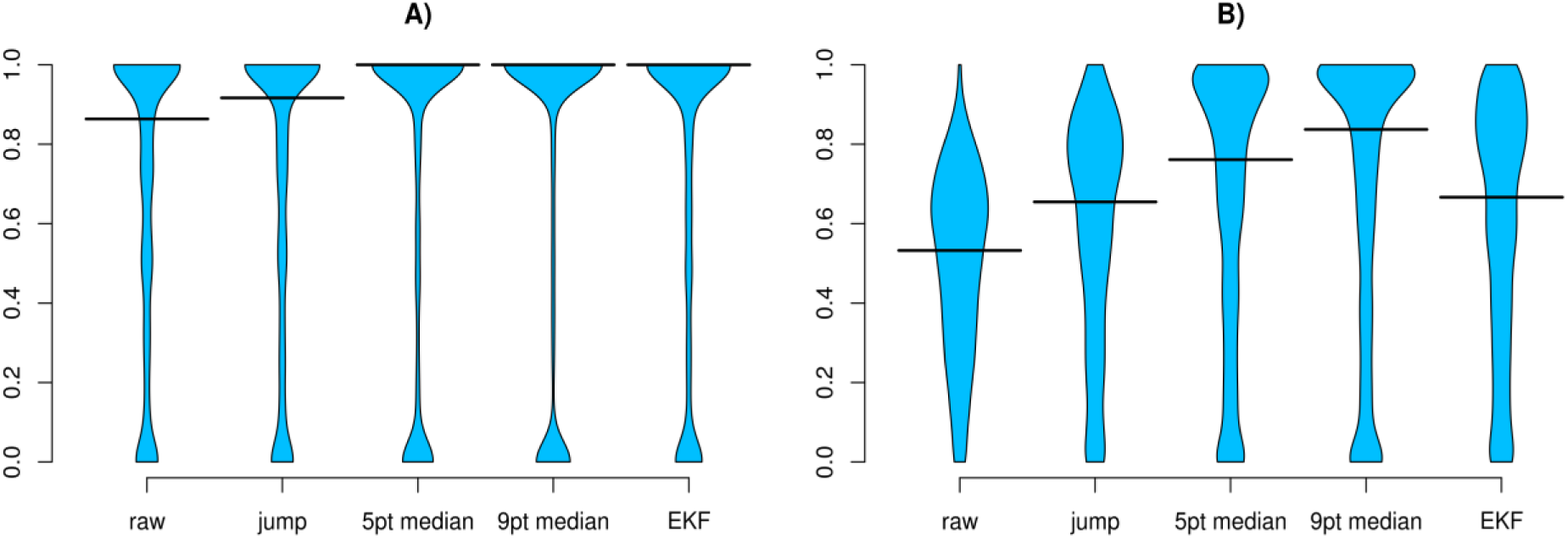
The proportion of points hitting the correct area as compared to reference for A) lying stalls B) Insentec feeding troughs

Figure 7 shows the density estimates for positioning data when the cows are feeding in the feeding fence on bottom edge of the barn. The contours depict 95% of all points during 24 hours of measurements. Filtering clearly increases the precision of the measurements, but does not improve the accuracy. The data for the troughs located in the center is located in the correct trough with quite low deviation. However, the feeders at the left and right edges are also located at the corners of the positioning area and show the greatest positioning error.

**Figure 7.**
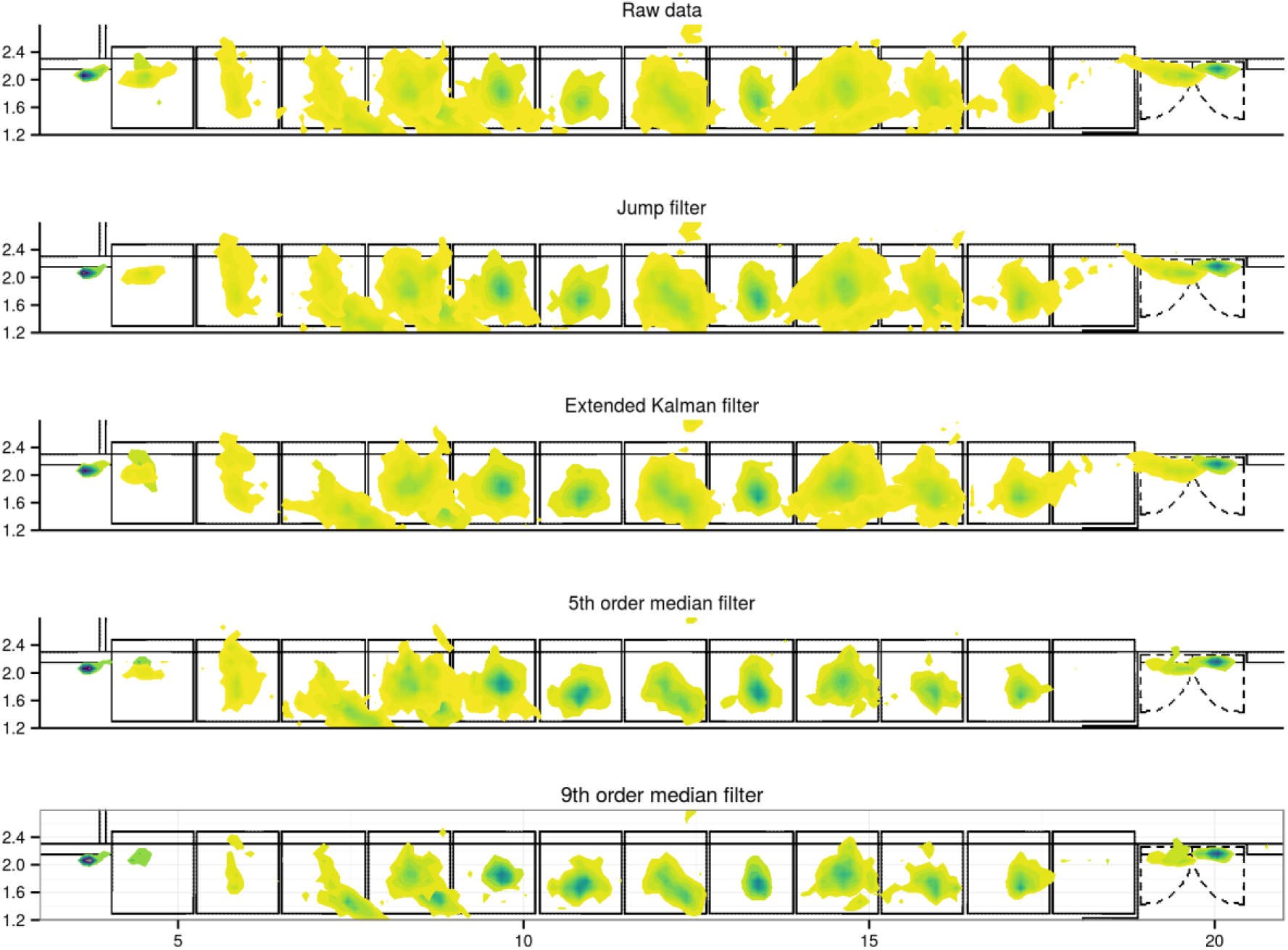
The effect of filtering on the positioning data while the cows are feeding on different locations of the feeding fence. The fence is located at the bottom edge of the barn and the boxes are the individual Insentec feeding troughs.

Figure 8 shows a closer look at the density estimates of the 9th order median filtered data for the troughs at the left corner, center and right corner relative to the center of the trough. The data for the left and right corner is not located inside the correct troughs, but distance to the feeding fence (y-axis) is very close to the true value.

**Figure 8.**
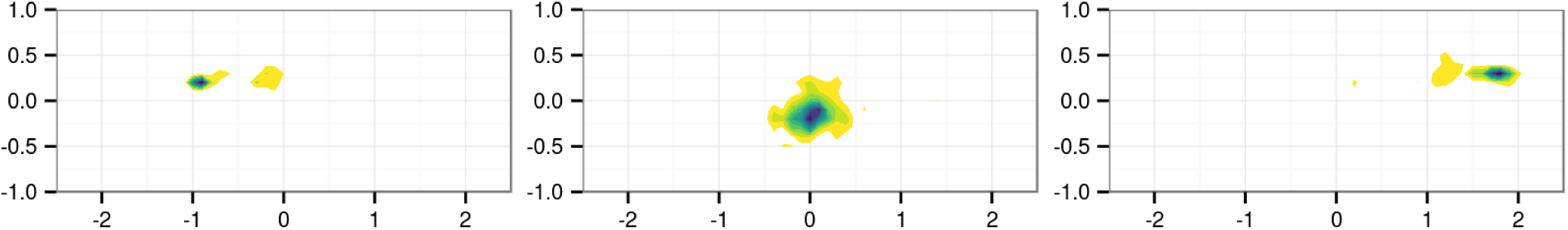
The density estimates of the positioning data for the troughs at the left corner, center and right corner relative to the center of the trough.

We also compared the trajectory of walking cows with raw data and different filters. Although we could not record the exact path of the cows we can make some general statements about the usefulness of different filters when the cows are walking. Figure 9 shows two examples of raw and filtered data of a walking cow. The left side shows a cow that is walking slowly (0.5 m/s) with no missing data and there is not a big difference between the performances of the different filters, although we can see that the median filters approximate the path as a straight line. The right side of Figure 9 shows a cow that is walking with higher speed 0.75 m/s and samples have been received only every 1.7s i.e. 50% of the data is missing. The 9^th^ order median filter clearly underestimates the path of the cow showing a route directly through the stalls whereas the extended Kalman filter makes a more realistic approximation of the cow’s path.

**Figure 9.**
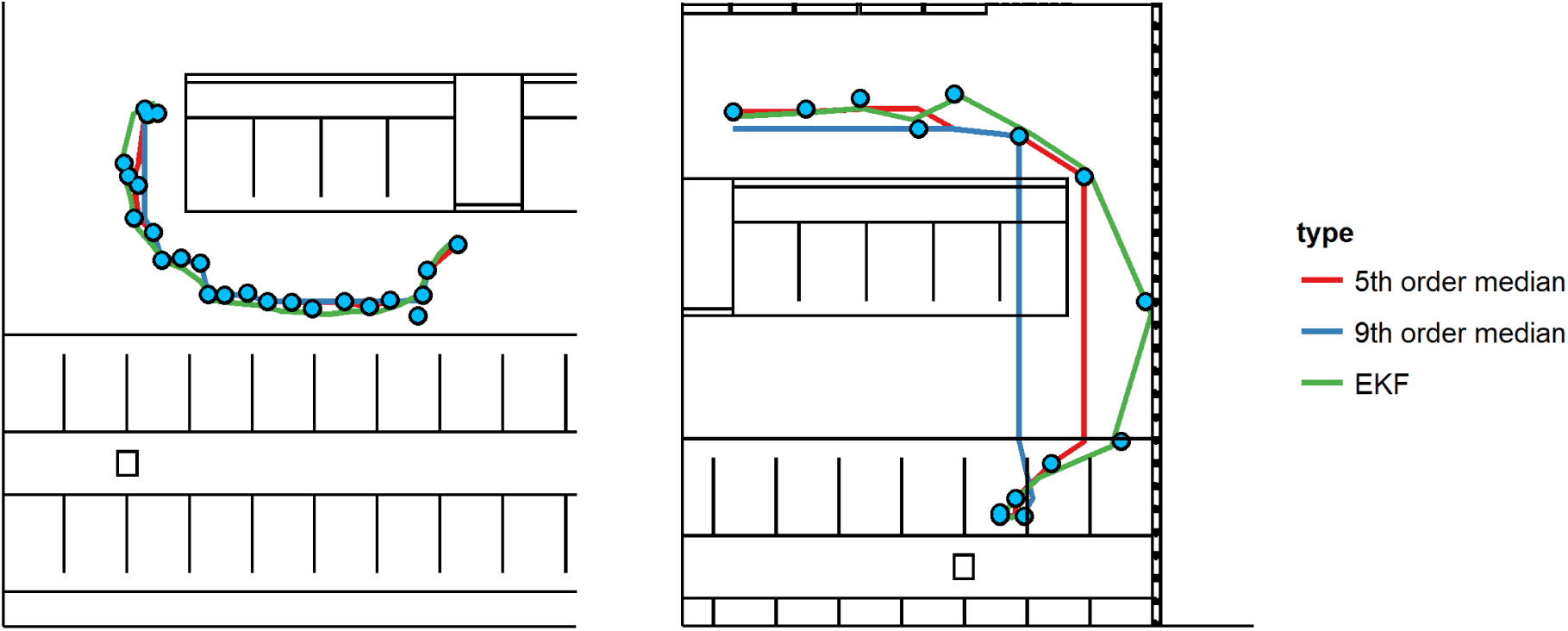
Two examples of raw (solid blue dots) and filtered data for a slowly moving cow with no missing data (left) and more quickly moving cow with 50% missing samples (right).

## 4. Discussion

We were able to improve the quality of the positioning data significantly using a heuristic jump filter with extended Kalman filter or median filter. The 9^th^ order median filter provided best estimates for cow position when the cows were not moving with median 100% of measurements located in correct stall and 84% in correct feeding trough when compared to the reference observations and measurements. The extended Kalman filter also improved the positioning accuracy significantly when compared to raw data and provides better of estimates of the trajectory of moving cows.

The choice of optimal filter therefore depends on the application where the measurement data will be used. If it is used to study the time that the cows spent in different areas it is better to use the median filter, however if the movement of the animals is of interest then the extended Kalman filter or lower order median filter is likely a better solution. Especially the extended Kalman filter needs to be combined with the jump filter to provide good estimates of position. An alternative implementation for the jump filter would be use validation gate (Bailey et al. 2006) in EKF based on measurement innovation.

The accuracy of the positioning systems varies depending on the location with relation to the sensors calculating the position from the tags. We got very accurate results for the feeding troughs in the center of the feeding fence with 95% of the data inside a circle with diameter of 50cm from the center of the trough. In contrast the median of the positioning data in the edges of the feeding fence and in the corners of the sensor coverage had a worst case error of close to 2 m. The accuracy of the setup in this study results is very similar to the one shown by Porto et al. (2014) also in a free stall barn. Similar findings about measuring the presence of a cow in a defined zone using GEA CowView have also been shown by Tullo et al 2016.

We encountered in average 58% missing samples during the trial. The reason for the missing data could be either lost signal or Ubisense built-in error rejection algorithm. However the results show that we got good estimates for cow position as compared to reference data after filtering and piecewise interpolation. Most of the data was missing when the cows were lying in the stalls and there were less samples missing when the cows were moving in the alleys. The effect of the missing data in measuring the correct path of the moving cows needs to be studied further.

The data from the positioning system could also be combined with data from an inertial measurement unit (IMU) using EKF to get more accurate positioning results. This approach has been shown to work well with UWB positioning systems and IMUs in human applications (Pulido Herrera 2009, DiFranco et al. 2014).

In the future combining positioning data with other sensors to automatically measure behaviors could yield higher prediction accuracy as different behaviors with similar movement patterns are more likely to appear in different functional areas e.g. feeding near the feeding area, drinking at the water trough and rumination at other areas.

## 5. Conclusions

The UWB positioning system provides accurate estimates of cow position inside a free stall after filtering. There was a large number of missing data which we were able to correct using interpolation. The choice of optimal filter depends on the intended use of the data and the effect of noise and missing data in the estimation of the correct path of moving cows needs to be studied in more detail.

## 6. Acknowledgements

The authors wish to thank the staff of Luke Maaninka research barn for technical assistance. We also thank students Maija Lappalainen and Sara Mykkänen for conducting the direct observations in the experiment.

## References

Backman, J., Oksanen, T., & Visala, A. (2010). Nonlinear model predictive trajectory control in tractor-trailer system for parallel guidance in agricultural field operations, in Proceecings of Agricontrol 2010, Kyoto, Japan, December 6-8, 2010, IFAC International Conference, 2010.

Backman, J., Oksanen, T., & Visala, A. (2013). Front Wheel Skidding Compensation System in Snow Ploughing, in Proceecings of Agricontrol 2013, Espoo, Finland, August 27-30, 2013, IFAC International Conference, 2013.

Bailey, Tim, Upcroft, Ben, & Durrant-Whyte, Hugh (2006). Validation Gating for Non-Linear Non-Gaussian Target Tracking. In Proceedings 9th International Conference on Information Fusion, 2006, IEEE, Florence, Italy.

Di Franco, C., Franchino, G. & Marinoni, M. (2014). Data fusion for relative localization of wireless mobile nodes, in Industrial Embedded Systems (SIES), 2014 9th IEEE International Symposium on, pp.58–65, 18-20 June 2014.

Eddelbuettel, D., & Sanderson. C. (2014). RcppArmadillo: Accelerating R with High-Performance C++ Linear Algebra. Computational Statistics & Data Analysis. 71, 1054–63.

Frondelius, L., Pastell, M., & Mononen, J. (2014). Validation of the TrackLab positioning system in a cow barn environment. In: Proceedings of Measuring Behavior 2014, Wageningen, the Netherlands, 27-29.8.2014, eds. Spink AJ, Loijens LWS, Woloszynowska-Fraser M and Noldus LPJJ. Wageningen University. 4 p.

Frondelius, L., Kajava, S., Lindeberg, H., Mononen, J., & Pastell, M. (2015). Measuring the effect of hoof lesions on cow’s walking, lying and eating behavior. In: Precision livestock farming ‘15, 7th European conference on precision livestock farming, Milan, Italy, 15-18 September’15/Eds. M. Guarino and D. Berckmans. p. 363–369.

Gygax, L., Neisen, G., & Bollhalder, H. (2007). Accuracy and validation of a radar-based automatic local position measurement system for tracking dairy cows in free-stall barns. Computers and Electronics in Agriculture. 56, 23–33.

Homer, E.M., Gao, Y., Meng, X., Dodson, A., Webb, R. & Garnsworthy, P.C. (2013). Technical note: A novel approach to the detection of estrus in dairy cows using ultra-wideband technology. Journal of Dairy Science, 96, 6529–6534.

Peter Kampstra (2008). Beanplot: A Boxplot Alternative for Visual Comparison of Distributions. Journal of Statistical Software, 28(1), 1–9.

Kalman, R.E., (1960). A new approach to linear filtering and prediction problems. Transactions of the ASME, Journal of Basic Engineering, 82, 34–45.

Lenain, R., Thuilot, B., Cariou, C. & Martinet, P. (2005). Model Predictive Control for Vehicle Guidance in Presence of Sliding: Application to Farm Vehicles Path Tracking. In Proceedings of the 2005 IEEE International Conference on Robotics and Automation, p. 885–890. ICRA, Barcelona, Spain.

Härdle, W. & Steiger, W. (1995) Optimal median smoothing, Applied Statistics, 44, 258–264.

Oberschätzl, R., Haidn, B., Peis, R., Kulpi, F., Völkl, C. 2015. Validation of automated processed position data for assessing dairy cow behaviour. Landtechnik 70(1), 2015, 3–8.

Oksanen, T., Linja, M., Visala, A. 2005. Low-cost positioning system for agricultural vehicles, IEEE International Symposium on Computational Intelligence in Robotics and Automation, Espoo, Finland

Piltaver, R., Cvetković, B. & Kaluza, B. 2015. Denoising Human-Motion Trajectories Captured with Ultra-Wideband Real-time Location System. Informatica 39 (2015) 311–322.

Porto, S.M.C., Arcidiacono, C. Giummarra, A., Anguzza, U., & Cascone, G. (2014). Localisation and identification performances of a real-time location system based on ultra wide band technology for monitoring and tracking dairy cow behaviour in a semi-open free-stall barn. Computers and Electronics in Agriculture, 108, 221–229.

Pulido Herrera, E. (2009). Improving data fusion in user positioning systems. Doctoral Thesis. Universitat Jaume I, Spain.

R Core Team. (2015). R: A Language and Environment for Statistical Computing. Vienna, Austria: R Foundation for Statistical Computing. www.R-project.org/.

E. Tullo, Fontana, I., Gottardo, D., Sloth, K.H. & Guarino, M. (2016). Validation of a commercial system for the continuous and automated monitoring of dairy cow activity. Journal of Dairy Science, 99(9), 7489–7494.

Tøgersen F.A., Skjøth F., Munksgaard L., & Højsgaard S. (2010). Wireless indoor tracking network based on Kalman filters with an application to monitor dairy cows. Computers and Electronics in Agriculture, 72, 119–126.

G.L. Smith; S.F. Schmidt & L.A. McGee (1962). Application of statistical filter theory to the optimal estimation of position and velocity on board a circumlunar vehicle. Technical Report R-135, National Aeronautics and Space Administration.

Veissier, I., Mialon, M-M., & Sloth, K.H. (2017) Early modification of the circadian organization of cow activity in relation to disease or estrus. Journal of Dairy Science, 100(5), 3969–3974

Wickham, H. (2009). Ggplot2: Elegant Graphics for Data Analysis. New York, USA: Springer.

